# Information use in foraging flocks of songbirds - no evidence for social transmission of patch quality

**DOI:** 10.1101/784777

**Authors:** F. Hillemann, E. F. Cole, B. C. Sheldon, D. R. Farine

**Affiliations:** Edward Grey Institute of Field Ornithology, Department of Zoology, University of Oxford, Oxford, U.K.; Department of Collective Behaviour, Max Planck Institute of Animal Behavior, Konstanz, Germany; Centre for the Advanced Study of Collective Behaviour, University of Konstanz, Konstanz, Germany; Department of Biology, University of Konstanz, Konstanz, Germany

**Keywords:** collective animal behaviour, economic decision-making, group foraging, local enhancement, Paridae, social information, social network

## Abstract

Animals use behavioural cues from others to make decisions in a variety of contexts. There is growing evidence, from a range of taxa, that information about the locations of food patches can spread through a population via social connections. However, it is not known whether information about the quality of potential food sources transmits similarly. We studied foraging behaviour in a population of wild songbirds with known social associations, and tested whether flock members use social information about the profitability of patches to inform their foraging decisions. We provided artificial patches (ephemeral bird feeders) that appeared identical but were either profitable (contained food) or unprofitable (contained no food). If information about patch profitability spreads via social associations, we predicted that empty feeders would only be sampled by individuals that are less connected to each other than expected by chance. In contrast, we found that individuals recorded at empty feeders were more closely associated with each other than predicted by a null model simulating random arrival of individuals, mirroring pattern of increased connectedness among individuals recorded at full feeders. We then simulated arrival under network-based diffusion of information, and demonstrate that the observed pattern at both full and empty feeders matches predictions derived from this post-hoc model. Our results suggest that foraging songbirds only use social cues about the location of potential food sources, but not their profitability. These findings agree with the hypothesis that individuals balance the relative economic costs of using different information, where the costs of personally sampling a patch upon arrival is low relative to the cost of searching for patches. This study extends previous work on information spread through avian social networks, by suggesting important links between how animals use information at different stages of the acquisition process and the emerging population-level patterns of patch use.

## INTRODUCTION

Animals can learn about their environment by observing others. Socially-acquired information involves any information gained through social processes, whether extracted from actively produced signals, or (inadvertent) cues provided by others’ behaviour and its consequences (Danchin et al. 2004; Smidt et al. 2010). Use of social information has been demonstrated in a wide variety of contexts including habitat choice (Forsman et al. 1998), assessing predation risk (Dawson & Chittka 2014), finding food (Tóth et al. 2017), and learning which food to avoid (Van de Waal et al. 2013; Thorogood et al. 2018). Social information about the availability of food is particularly valuable in unpredictable ephemeral environments (Baude et al. 2008; Boyd et al. 2016), when acquiring personal information about the environment can be costly in terms of time and energy (Valone & Templeton 2002; Kendal et al. 2005). Models assuming perfect information-sharing among group members suggest that groups have a higher searching efficiency than individuals, resulting in improved patch-finding rate with increased group size, and that groups are more efficient in patch assessment, reducing the time spent in unprofitable patches (e.g., Clark & Mangel 1986; Ranta et al. 1993). However, information acquired through personal experience can be more accurate, or relevant, than social information. For example, individuals can differ in their preferences or needs with regard to resource quality, and observing one individual choose, or reject, a patch could yield incorrect information to the observer. Using social information could also represent an ecological trap (Schlaepfer et al. 2002)—if many patches are unprofitable, it could cause individuals to spend more time finding food. Thus, foragers faced with a trade-off between gathering more accurate (personal) information or cheaper (social) information, should use social information when personal information cannot be gathered reliably at a low cost, and otherwise rely on personal experience (Kendal et al. 2005; Galef 2009; Czaczkes et al. 2019).

Studies across a range of taxa have found that information about the location of food patches spreads through a population via social connections, i.e. individuals became more likely to discover novel resources that are widely dispersed throughout the landscape when they were socially connected to knowledgeable conspecifics or heterospecifics (e.g., Aplin et al. 2012; Webster et al. 2013; Farine et al. 2015a; Jones et al. 2017; Tóth et al. 2017). Further, birds foraging in mixed-species flocks have been found to move between clumped resources towards patches with the largest proportion of the flock (Farine et al. 2014). Together, these results suggest that songbirds use social information both when finding patches, and when deciding where to forage within a patch. However, it is not always clear what cues animals are responding to, or whether information is being transmitted between individuals, because different social and asocial mechanisms can lead to similar pattern of apparent spread of information.

Identifying what information individuals acquire and use is important for understanding social influences on foraging (Galef & Giraldeau 2001). For example, it may not be immediately obvious whether individuals arriving at a new resource discovered the patch together (i.e., simultaneously), or whether information is being transferred from individuals that are knowledgeable of the availability of resources to naïve individuals. To differentiate the effect of collective movement from social transmission of information about the patch location, some studies have compared the order in which individuals arrived at foraging patches with the order of arrival at control patches (Atton et al. 2012), while others adjusted the analysis to account for the possibility that no information about the patch location had been transferred between individuals that arrived within a short period of time (e.g., Aplin et al. 2012). Similarly, individuals might use personal or social information to assess the quality or profitability of foraging patches. There is mixed evidence on whether individuals determine foraging patch quality from the behaviour of others; the use of social information in this context has been reported for European starlings, *Sturnus vulagaris* (Templeton & Giraldeau 1995), red crossbills, *Loxia curvirostra* (Smith et al. 1999), and three-spined sticklebacks, *Gasterosteus aculeatus* (Coolen et al. 2003), but there was no evidence this was the case in studies of European blackbirds, *Turdus merula* (Smith et al. 2001), or nine-spined sticklebacks, *Pungitius pungitius* (Coolen et al. 2003). Further, it is not known whether information about patch quality transmits via social connections, in a similar fashion to the way information about the location of foraging patches has been shown to spread through populations.

Here, we study foraging behaviour in mixed-species flocks of tits to test whether individuals use social information when assessing foraging patches. Tits and other species from the Paridae family provide a good system for information diffusion experiments, because they readily use social information in foraging contexts, from finding food (e.g., Sasvári1979; Farine et al. 2015b; Firth et al. 2016; Hillemann et al. 2019a), to learning innovative foraging techniques (Aplin et al. 2013, 2015) and avoidance of unpalatable prey (Thorogood et al. 2018). Further, it has been shown that information does not spread between individuals at random but can be predicted by their connections in the population-level social network (Aplin et al. 2012; Farine et al. 2015a; Firth et al. 2016). Yet, it is not clear whether social cues are also used to assess the quality, or profitability, of foraging patches.

We expand previous studies on social foraging in songbirds, by contrasting pattern of information spread for patches that varied in profitability, to test whether songbirds use social information about whether a patch contains food or not. We provided artificial patches (bird feeders) that appeared identical but were either empty or stocked with seed, and calculated the connectedness of individuals arriving at each feeder using the edge strength from independently-recorded social association network data. We then compared the order in which birds discovered these feeders with a null model simulating random arrival of birds, to test two distinct hypotheses.

The first hypothesis is that, additionally to information about patch location, birds also acquire social information about patch profitability by attending to the behaviour of others. If individuals gain information from their close associates that the patch was not rewarding (here, absence of food), we expect them to not sample the patch themselves. Thus, under this hypothesis, we predict that individuals sampling empty feeders will be less connected to those that had previously visited the feeder (“knowledgeable individuals”, hereafter), than would be expected by the random arrival model, because closely-associated individuals are more likely to learn from each other (e.g., Coussi-Korbel & Fragaszy 1995). At profitable feeders, on the other hand, we expect that arriving birds should be more connected to the knowledgeable individuals in their social network than predicted by the null model.

The alternative hypothesis is that birds acquire only information about the location of potential resources, but not their quality. Individuals may not obtain the relevant social information to assess whether a patch is profitable, or they may not use it. Under this hypothesis, individuals discovering a potential patch are therefore expected to sample it regardless of its quality. Thus, we expect that the connectedness of arriving individuals should be higher than predicted by the random arrival model, regardless of whether the feeder is full or empty. In other words, the results should reflect the same social effects on patch discovery that have been shown in other studies (Aplin et al. 2012; Farine et al. 2015a; Firth et al. 2016).

## METHODS

### Study system

We studied a population of individually tagged songbirds in Wytham Woods in Oxfordshire, UK (51°46’ N, 1°20’ W), consisting of blue tits (*Cyanistes caeruleus*), great tits (*Parus major)*, marsh tits (*Poecile palustris)*, and Eurasian nuthatches (*Sitta europaea*). Tits in this population move and forage in mixed-species flocks during the non-breeding season (Hinde 1952; Farine et. al 2012, 2014, 2015a), with blue tits and great tits being the most abundant species, that together accounted for more than 90 % of the bird recorded in this study. Birds were each fitted with a unique British Trust for Ornithology (BTO) metal leg ring as nestlings or as adults, either when caught at nest boxes during the breeding season or in mist nets during the winter. Additionally, birds were fitted with a plastic leg ring carrying a passive integrated transponder (PIT) tag (IB Technologies, UK), that can be read by radio-frequency identification (RFID) antennae.

### Social association data

Following the same protocol as previous studies on our system (e.g., Aplin et al. 2012, Farine et al. 2015b, Hillemann et al. 2019b), we quantified social associations of individuals based on co-occurrence in foraging groups. Feeder visits (time and location) by PIT-tagged birds were recorded at automated feeding stations with RFID antennae (Dorset ID, Netherlands), that were spread in a grid of 250 x 250 m across the study area (56 sampling sites in the first season, and 54 sites in second season). These feeding stations were accessible for a two-day sampling periods per week over multiple consecutive weeks in January and February (first season: eight sampling periods, second season: six sampling periods), thus providing a data stream detailing the spatio-temporal distribution of birds. To identify gathering events, or flocks, we used a Gaussian mixture model (GMM) that was designed to detect periods of high activity (Psorakis et al. 2012, 2015). The GMM approach is a clustering algorithm that can identify groups of different sizes and temporal resolution, without the need to set arbitrary time resolution parameter (e.g., fixed time windows). For detecting groups, we used the gmmevents function in the asnipe package (Farine 2013) in R (v. 3.4.4; R Development Core Team 2018).

To calculate a “local” network for each experimental site, we used subsets of the population-scale social association data. For each trial, we included data collected at the four logging stations closest to the experimental site, and during the four two-day sampling periods closest to the feeder discovery trial. GMM models were applied to the data from each feeder on each day, and we aggregated data from all of the flocks into one group-by-individual matrix for each of the local networks. Social association was assumed based on co-presence in the same gathering event, and association strengths for each dyad was calculated as the simple ratio index, which ranges from 0 (never recorded in the same foraging flock) to 1 (always recorded in the same foraging flock). Social networks were constructed using the asnipe R package (Farine 2013).

### Patch-discovery experiment

We conducted feeder discovery trials at 18 different sites (10 sites in February - March 2016, 8 sites in February - March 2017). Experimental sites in the same year were separated by at least 350 meters, and 88% of the individuals (N = 538) recorded throughout the experiment were either only ever recorded at one experimental site, or at only one site in each of the two years (Figure S1 in the Supplementary Material). We implemented a fully balanced design, presenting both treatments (full and empty feeder) at each site. This design allowed us to balance potentially confounding factors such as variation in the local density of birds and habitat structure. The order of trials was pseudo-randomised, and we left a minimum of 3 days between consecutive trials at a given site (average ± standard deviation: 10.3 ± 7.2 day). We deployed feeders after sunset on the evening before the trial to ensure natural patch discovery the next morning. The feeders were either stocked with unhusked sunflower seeds, or completely empty, and covered with an opaque tube (grey PVC pipe). Two access holes to the feeder were fitted with RFID antennae (scanning rate: 16 Hz), recording the identity of all PIT-tagged birds inspecting the feeder or taking a seed (one record per individual in each 15-s interval). Discovery trials lasted one day, and feeders were removed immediately after the end of the trial. Data from full-feeder trials were previously published in Hillemann et al. 2019a, in the context of analysing diurnal foraging patterns.

### Modelling the spread of information about food availability

For each trial, we recorded the order of arrival of k individuals, with k being the total number of tagged birds that discovered the patch. Birds were classified as knowledgeable after their first visit was detected at the patch. For each arriving bird, we calculated its mean edge strength to knowledgeable individuals, using the independently collected social association data. We then compared the observed pattern of patch discovery to a null model simulating random arrival of individuals by selecting k individuals at random from the network. A random discovery pattern is expected if individuals are not using social cues, but instead discover and sample the food source independently. We ran 1000 repetitions of this simulation, and compared the observed social connectivity between knowledgeable individuals to the model prediction.

Since birds that were recorded at the feeders were more connected than predicted by the null model in both treatments (full and empty feeders), we then developed a network-derived transmission model, the ‘affiliative model’, to test whether the observed pattern of arrival at the feeders could be explained by social associations. We randomly selected one individual to make the initial discovery, then randomly chose one of its associates as the second individual to arrive at the feeder. For each subsequently arriving bird, we randomly chose an individual from the growing number of associates to knowledgeable individuals. We repeated this process until k individuals were selected. Values obtained from 1000 repetitions of this simulation were again qualitatively compared to the observed pattern. Running these random-arrival simulations and network-derived simulations allowed us to quantify how the strength of connections between new discoverers and knowledgeable birds was expected to increase as the pool of information grows in a local population, under different mechanisms of information use.

### Ethical Note

This work was conducted as part of a large ongoing research project on Parid social behaviour at Wytham Woods. All work was subject to review by the local ethical review committee of the Department of Zoology, University of Oxford. Birds were caught, ringed, and PIT-tagged by experienced ringers under BTO licences.

## RESULTS

All of the full feeders (n=18 out of 18 trials) and all but three of the empty feeders (n=15 out of 18 trials) were discovered. Compared to full feeders, empty feeders were discovered by fewer individuals, and were visited less often throughout the length of the trial (Fig. 1). Empty feeders were only visited by few individuals in the early-morning hours, whereas birds accumulated at full feeders throughout the morning (Fig. 1a). The average number of birds visiting empty feeders was 10.2 ± 2.3 standard error mean (SEM; n=15), compared to 36.1 ± 6.8 SEM at full feeders (n = 18). Empty feeders were on average visited 25.5 ± 6.2 SEM times (per capita visit rate: 2.3 ± 0.3), whereas the average number of visits to full feeders was 645.2 ± 146.0 SEM, with per capita visit rates of 39.8 ± 5.3 SEM visits per individual per trial.

**Figure 1:**
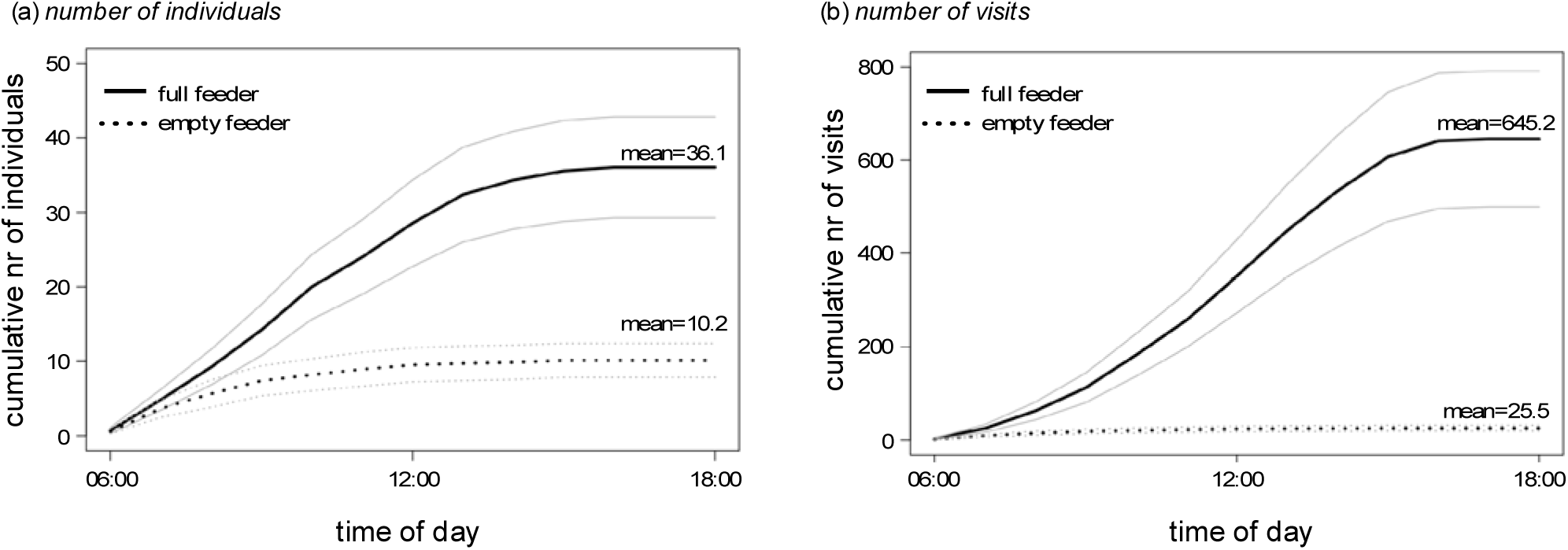
Cumulative number of (a) individuals visiting, and (b) feeder visits at full (continuous line) and empty feeders (dashed line) per hour of the day. Bold lines represent mean values across all trials in which the feeder was visited by at least one individual (18 full-feeder discovery trials and 15 empty-feeder trials, out of 18 trials conducted for each condition), and thin lines show standard errors of the mean. The total number of individuals visiting full feeders was on average 3.5-times higher than the number of birds visiting empty feeders, and full feeders were on average visited 25-times more than empty feeders.

At empty feeders, knowledgeable individuals were more connected to each other than predicted by the null model of random arrival (Fig. 2). For both feeder types, new individuals arriving at a feeder had a much higher edge strength to those birds that had already experienced the profitability of the patch, relative to the same number of individuals drawn randomly from the local population. The affiliative model, which simulates network-derived pattern of discovery, more accurately predicted the observed connections among individuals arriving at either feeder (empty or full), with average values of edge strength to knowledgeable individuals falling within the 95% range of the values obtained from the affiliative model simulations.

**Figure 2:**
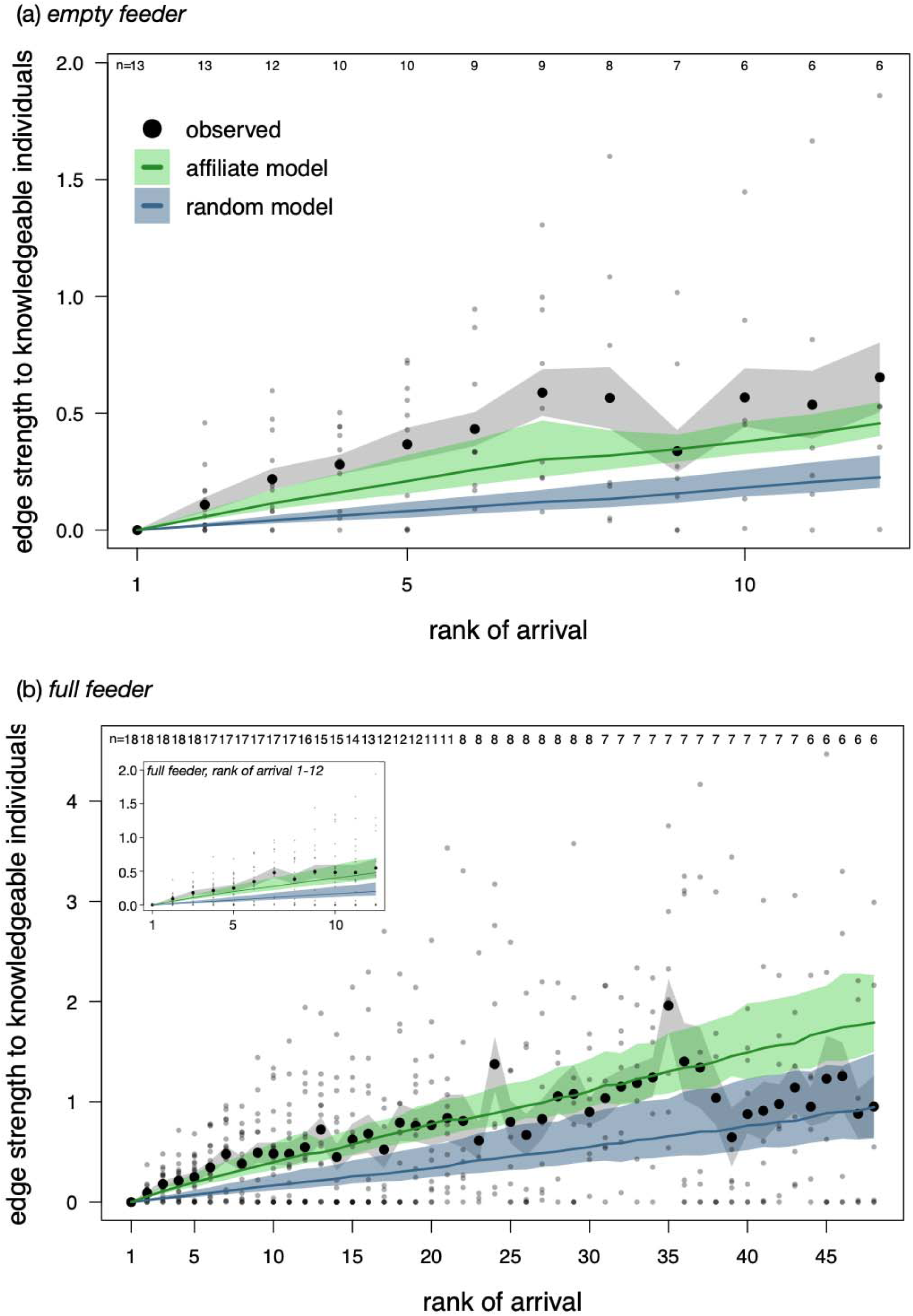
Mean edge strength of arriving individuals to birds that have already visited (a) empty feeders, or (b) full feeders. Small grey dots show observed data from single trials, larger black dots show mean values over all trials, and grey-shaded areas represent the mean ± SEM of the observed data (number of trials is given at the top margin of the plot). Only trials in which at least two individuals visited the feeder were included, and the plots only show up to the k-th rank of arrival that was observed in at least six trials (rank 1-12 for empty feeder trials, rank 1-48 for full feeder trials; the smaller frame in (b) shows only ranks 1-12 for full feeder trials). Blue and green and blue lines are means of expected values according to 1000 simulations of the random and affiliate model, and shaded areas represent the 95 % ranges of the simulated data, respectively. Perfect transmission of social information about patch quality would result in a straight line along 0 for empty feeders (i.e., only unconnected individuals visiting the unprofitable patch). Instead, individuals’ arrival to a feeder - empty or full- was strongly predicted by social connections suggesting that only information about the location of a potential resource is being transmitted socially, and not information about the quality of the patch. The drop in average connectedness to knowledgeable individuals, after the arrival of ca 35 to 40 individuals, indicates new independent discovery events. The average number of birds visiting full feeders was 36.1 ± 6.8 (mean ± SEM, n = 18), and 10.2 ± 2.3 at empty feeders (mean ± SEM, n = 15).

## DISCUSSION

By providing artificial patches that differed in their profitability, we found that non-rewarding patches were visited less often, and by fewer individuals, than patches that contained food. However, our results do not support the hypothesis that individuals acquire and use social information to assess the profitability of potential patches. Our prediction was, that empty patches should have been sampled only by individuals that are less connected to each other than the same number of randomly chosen individuals from the local population. Instead, we found that individuals arriving at empty feeders were more connected to each other than expected by chance. These results closely matched the patterns observed at independent trials presenting full feeders. Thus, we find no evidence that tits use social information to assess the profitability, or quality, of foraging patches.

From our data, we cannot conclude whether social information about patches not being profitable is (i) not available to other foragers, (ii) is not collected, or (iii) if foraging individuals actively forgo social information about patch quality (i.e., information is available but not used). We expect that individuals should observe different behavioural cues from others experiencing a full versus an empty patch (e.g., leaving with or without carrying a seed in their bill). However, individuals may not have had the opportunity to carefully observe potentially subtle differences in others’ behaviour, but instead sample personal experience about the patch immediately upon arrival. Alternatively, individuals may ignore the presented information (i.e., observe others return after unsuccessful feeder visits) in favour of sampling potentially more relevant and reliable personal information.

Using personal information to evaluate the quality of patches supports existing theory on economic decisions relating to social information use strategies. A number of theoretical models and empirical studies have suggested that personal and social information are not weighed equally, and that the relative use of social and personal information should depend upon the costs involved with acquiring and using each (e.g., Valone &Templeton 2002; Kendal et al. 2005; Hillemann et al. 2019a). Whereas personal exploration and trial-and-error sampling of the environment can be relatively costly, the cost of gathering personal information about the quality of a patch upon arrival is much smaller (Kendal et al. 2005; Galef 2009). Our results align with the predictions of this theory. Wintering songbirds seem to balance the costs of using personal versus social information during foraging by using social information to find potential resource locations, but not when evaluating the quality of a patch (which suggests that they then rely on personal information).

An economic view on when animals should use social versus personal information is consistent with recent findings about conformist transmission in our population (Aplin et al. 2015a). Conformity occurs where individuals ignore personal information, or forgo sampling it, and instead match their choices with the majority behaviour in the population (e.g., Kendal et al. 2004; Van de Waal et a. 2013; Aplin et al. 2015b). Behavioural conformism, such as visiting a patch together even if it is unprofitable, has been suggested to facilitate greater cohesion at the pair-level (Firth et al. 2016), and group-level (Aplin et al. 2014). The potential loss of contact with flock members and loss of anti-predator benefits associated with social foraging (e.g., Lima 1995) could select for social information use strategies that contribute to maintaining group cohesion above other factors. Formation of and moving together in foraging flocks is important for small songbirds such as tits. Further, during the winter, tits depend on dispersed, clumped food resources such as beech mast (the seeds of beech, *Fagus sylvatica*; Hinde 1952). Using social information to find potentially rich resources significantly reduces the time to patch discovery (e.g., Hillemann et al. 2019a), and this benefit may outweigh the costs of occasionally getting incorrect information.

Although birds appeared to use information similarly when discovering full versus empty feeders, we found that the number of individuals that visited feeders in the different conditions different consistently. On average, 3.5-times more individuals arrived at full feeders than at empty feeders. The differences in the number of arrivals in the two conditions highlights how beneficial group-level properties—in this case having more individuals discover profitable than non-profitable food sources—can emerge as a result of the strong social reinforcement in patch use by species such as tits (Aplin et al. 2014; Farine et al. 2014). Our data also suggest that birds visited empty patches because they acquired “wrong” information from observing others, demonstrating that social information use can also be costly, or represent an ecological trap (Schlaepfer et al. 2002), hindering effective foraging by causing large numbers of birds in a population to sample potentially suboptimal patches before acquiring personal information. However, we observed that individuals departed from a patch once they updated their personal information, thus reducing the time available for others to also acquire incorrect social information. We also know that birds in our population are unlikely to become trapped by incorrect information. A previous study of information transfer in our population that used puzzle-boxes that gave low rewards for common solutions and high rewards for uncommon solutions found that populations did not get trapped by their conformist choices or their personal information. Instead, populations could switch to high reward solutions as a result of the dynamic structure of the social network they live in (Aplin et al. 2017). Work in fish has suggested that social interactions among individuals can facilitate ‘collective sensing’, or the fine-scale tracking of environmental features without requiring any group-level control (Berdahl et al. 2012). How social information use among individuals translates to group-level, or population-level, movement and patch use decisions is an exciting area for future research.

Our study extends previous work on information use in a foraging context (e.g., Aplin et al. 2012) to test how individuals use social cues at different stages when navigating their environment. We build on previous approaches using populations with known social structure to track the acquisition of social information (Duboscq et al. 2016), such as studies using network-based diffusion analysis (Franz & Nunn 2009, Hoppitt et al. 2010). In our study, we used simulations to test for the expected patterns of connections between new arrivals and previous discoverers for each experimental trial. These models allowed us to evaluate alternative hypotheses about social information use at different steps in the acquisition process. Although very simple, our simulation of information spread through the network accurately characterised how information about food patches spread through the local social networks. Our study adds to the growing evidence for the importance of social information, and suggests that flexible information use and social learning strategies may prevent ecological traps, where suboptimal behaviour spreads through the population, even if individuals sometimes acquire wrong information.

## ACKNOWLEDGEMENTS

We are very grateful to Sara Keen for involvement in the data collection. We thank all Wytham fieldworkers for helping with the catching and marking of birds, and particularly Keith McMahon for his role in collecting the social association data. FH was funded by Natural Environment Research Council studentship award (reference: 1654580), DRF was funded by the Max Planck Society and the DFG Centre of Excellence 2117 ‘Centre for the Advanced Study of Collective Behaviour’ (ID: 422037984).

## SUPPLEMENTARY MATERIAL

### Independence of trials

**Figure S1.**
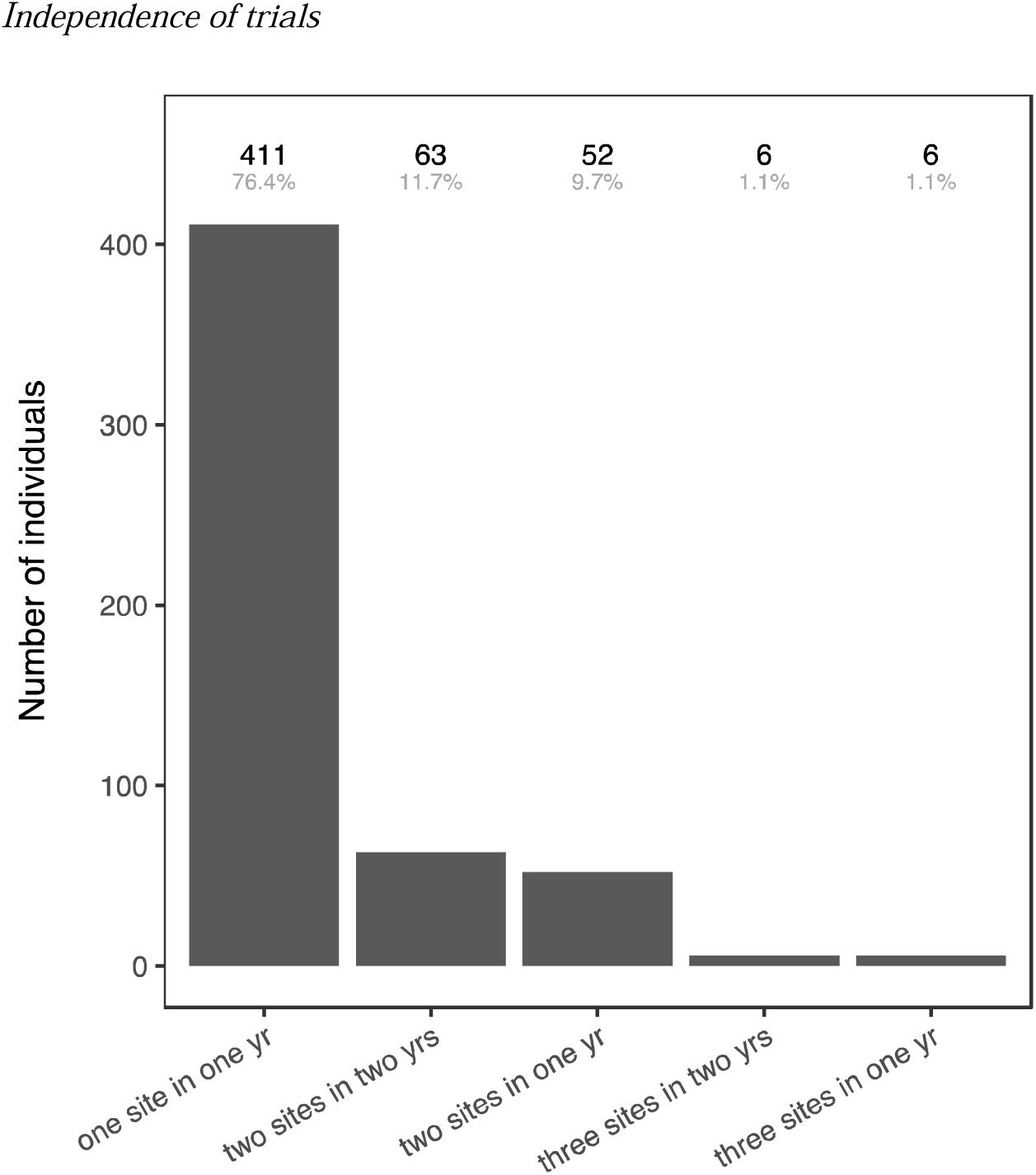
Number of individuals that were recorded at one or multiple experimental sites. Discovery trials with artificial patches that varied in their profitability (full and empty bird feeders) were conducted at 18 unique sites spread across the study area, and in two consecutive winters (10 sites in early 2016 and 8 sites in early 2017). Within the same year, experimental sites were separated by at least 350 meters. Represented are the number of individuals that were recorded at only one site, or at multiple sites (because they were recorded at neighbouring sites either in different years, or within the same year). Black numbers at the upper margin of the plot give the total number of individuals recorded per category, percentages are given in grey. Of all the 538 birds included in our study, 474 individuals (or 88%) were either only ever recorded at one experimental site throughout the study, or at only one site in each of the two years.

